# Enhanced inhibition of influenza virus infection by peptide-noble metal nanoparticle conjugates

**DOI:** 10.1101/324939

**Authors:** Zaid K. Alghrair, David G. Fernig, Bahram Ebrahimi

**Author notes:** Address for correspondence Department of Functional and Comparative Genomics, Institute of Integrative Biology, Biosciences Building, Crown Street, University of Liverpool, Liverpool, L69 7ZB, U.K.

## Abstract

Influenza virus is a major medical and veterinary health concern and causes global pandemics. The peptide ‘FluPep’ is an established inhibitor of influenza virus infectivity in model systems. We have explored the potential for FluPep functionalised noble metal nanoparticle to enhance the antiviral activity of the peptide Flupep and determined their potential for the delivery of FluPep. The FluPep ligand designed here is FluPep extended at its N-terminus with the sequence CVVVTAAA-, to allow its incorporation into a mix matrix ligand shell of a peptidol and an alkanethiol ethyleneglycol comprising 70% H-CVVVTol and 30% HS(CH_2_)_11_(ethyleneglycol)_4_ (both mole/mole). Gold and silver nanoparticles (~10 nm diameter) prepared with up to 5% (mole/mole) FluPep ligand contained in the mixture of mix-matrix peptide ligands remained as stable as the control mix-matrix coated nanoparticles against ligand exchange with dithiothreitol. FluPep ligand was found to inhibit viral plaque formation in canine MDCK cells (IC_50_ 2.1 nM), but was less potent than FluPep itself (IC_50_ 140 pM). FluPep ligand functonalised nanoparticles retained antiviral activity in the plaque assay. Moreover, at low grafting densities (where nanoparticles incorporate ~1 FluPep ligand, the antiviral potency in terms of FluPep ligand concentration was enhanced significantly for gold and silver nanoparticles (IC_50_ ~8-fold and ~3-fold lower, respectively). At higher grafting density the potency relative to free FluPep ligand concentration decreased. The data demonstrate that conjugation of FluPep to gold and silver nanoparticles enhances its antiviral potency; the antimicrobial activity of silver ions may enable the design of even more potent anti-microbial inhibitors.

## Introduction

Influenza type A (‘flu) virus is a major health concern for humans and livestock animals. The mode of transmission is by the respiratory route. ‘Flu infection occurs seasonally and can cause global pandemics. The most recent ‘flu pandemic was the 2009 H1N1 subtype swine ‘flu, which resulted in more than 18,000 deaths worldwide (Pasricha et al., 2013). The treatment of ‘flu infections is difficult, because the virus has an RNA segmented genome, which has a high potential to recombine and create new strains through a mechanism termed re-assortment (Treanor, 2004). Other potential risks to human populations are the zoonotic avian (bird) and porcine (swine) ‘flu viruses. Vaccination remains the most effective means to prevent and control infection (Grohskopf et al., 2011). However, the lead time to vaccine production is around 9 months, efficacy is not always complete, only a fraction of the human population is vaccinated and although some vaccines have been trialled against avian ‘flu, farm animals in general are not routinely vaccinated on a global scale. There is, therefore, the need for drugs to treat ‘flu.

Currently there are two common types of anti-influenza drugs, based on their mechanism of action. The first class are neuraminidase inhibitors such as Oseltamivir (Tamiflu). The second class are virus ion-channel blockers, such as Amantadine (Symetrel). The effectiveness of Tamiflu has been questioned (Cohen, 2014), and in any case emerging resistance of the influenza virus is leading to reduced effectiveness. The promise of peptide-based antiviral drugs is established by the Food and Drug Administration’s (USA) approval of Enfuvirtidie against HIV (Cooper and Lange, 2004). The mode of action of antiviral peptides falls into three categories which are: (i) entry blocker peptides, which inhibit viral attachment and virus-cell membrane fusion; (ii) viral envelope targeting peptides, which disrupt the viral envelope; (iii) viral polymerase peptides, which inhibit replication of virus genome inside the infected cell by interacting with viral polymerase (Skalickova et al., 2015). *In vitro*, several viral diseases have been treated with peptides such as HIV (Lee et al., 2014), (Pessi et al., 2009), hepatitis C (Abe et al., 2007), herpes simplex (Jenssen, 2005), (Jaishankar et al., 2015), and influenza virus (Albericio and Kruger, 2012, She et al., 2008, Nicol et al., 2012).

Infectivity of influenza A viruses, including the H1N1 subtype, is strongly inhibited by a peptide called FluPep (Nicol et al., 2012). FluPep was originally identified as a sequence in tyrosine kinase inhibitor peptide (Tkip), thought to act as a mimic of the suppressor of cytokine signalling (SOCS) protein (Ahmed et al., 2009). However, FluPep’s antiviral activity doesn’t depend on blocking suppressor of cytokine signalling, which is intracellular, but instead this peptide appears to exert its antiviral activity from the outside of the cell. Thus, addition of FluPep to cells in culture prevents infection by ‘flu virus, as does intranasal delivery of the peptide in a murine model of human ‘flu (Nicol et al., 2012).

Noble metal nanoparticles possess a strong plasmon absorbance, which allows detection at very low levels, using a range of approaches, from absorbance (Free et al., 2012, Haiss et al., 2007a) to photothermal microscopy (Berciaud et al., 2004) and various extensions of the latter (Lasne et al., 2006, Nieves et al., 2015). Noble metal nanoparticles can be passivated and functionalised with biomolecules such that they possess the biological selectivity and specificity of the grafted biological functional entity (Duchesne et al., 2012), which includes peptides (Maus et al., 2010). Presentation of a functional peptide by means of a nanoparticle has a number of advantages. Thus, nanoparticle conjugation may enhance the solubility of the peptide, as well as enhance its biological activity of the peptide, for example, due of multivalent functionalisation of the nanoparticles. In addition, noble metals such as silver possesses antimicrobial activities (Galdiero et al., 2011). Thus, noble metal nanoparticles are potentially useful as both functional probes for antiviral peptides and as therapeutic delivery platforms. We have, therefore, synthesized gold and silver nanoparticles functionalised with FluPep and analysed the anti-flu activity of the nanoparticle-FluPep ligand conjugates. The results demonstrate that the nanoparticle-FluPep ligand conjugates reduce the infectivity of influenza virus with greater antiviral activity than the free peptide, making this a viable tool for development of a peptide formulation that efficiently combats seasonal, pandemic, and zoonotic ‘flu Infections.

## Materials & methods

### Materials

#### Peptides and gold nanoparticles

Peptides FluPep WLVFFVIFYFFRRRKK, FluPep ligand CVVVTAAAWLVFFVIFYFFRRRKK and the ligand shell matrix peptidol CVVVT-ol were purchased from Peptide Protein Research (PPR Ltd, Hampshire, UK). The alkethiol ethylene glycol ligand, HS-EC_11_-EG_4_, was purchased from Prochimia (ProChimia Surface Sp. z o.o., Sopot, Poland). Gold nanoparticles of 9 nm diameter stabilized in citrate buffer were purchased from British Biocell (BBInternational Ltd, UK) and silver nanoparticles of ~10 nm diameter from nanoComposix Inc. (CA, USA). Nanosep filters 10 kDa cut off were from PALL (PALL Corp., Portsmouth, and Hants, UK). Uv-vis spectra (2 nm incremental steps) were measured using a SpectraMax Plus spectrophotometer (Molecular Devices, Wokingham, UK) and 384 well plates from Corning (Lowell, US) and the concentration of gold nanoparticles and of silver nanoparticles determined at 450 nm (Haiss et al., 2007b) and 392 nm (Paramelle et al., 2014), respectively.

#### Synthesis of FluPep functionalised nanoparticles

Mix matrix ligands 70:30 (mole:mole) CVVVT-ol: HS-(CH2)_11_-EG_4_-OH were prepared as described (Duchesne et al., 2008a) by first diluting 35 μL CVVVT-ol (4 mM DMSO:H_2_O) with 35 μL ddH_2_O and 6 μL HS-C_11_-EG_4_-OH (2 mM) with 6 μL EtOH and 18 μL ddH_2_O. Adding the two solutions together yielded a 2 mM ligand solution of 70% (mole/mole) CVVVT-ol and 30% (mole/mole) HS-C_11_-EG_4_-OH. The ligand mixture was added to 900 μL (gold or silver) nanoparticles and vortex-mixed. Once mixed, 100 μL of 10 × phosphate-buffered saline (PBS: 137 mM NaCl, 3 mM KCl l, 8 mM Na_2_HPO_4_, 15 mM KH_2_PO_4_) with Tween-20 (0.1 % v/v) pH 7.4 was added to gold nanoparticles (Duchesne et al., 2008a) and 10 × (100 mM NaNO_3_, 20 mM HEPES (Free et al., 2012) with Tween-20 (0.1 % v/v) pH 7.4 to silver nanoparticles (Free et al., 2012), vortex mixed and the (gold or silver) nanoparticles placed on a rotating wheel for 24 h. Nanoparticles were concentrated 10-fold using 10 kDa Nanosep centrifugal filters (PALL Corp., Portsmouth, Hants, UK). Samples were then centrifuged for 7 min at 10000 rpm (~12,000 x g) and the gold nanoparticles diluted with 1 × PBST (PBS 0.05% (v/v) Tween-20) and silver nanoparticles with 1x (100 mM NaNO_3_, 20 mM HEPES). The nanoparticles were then further separated from excess ligands by applying them (100 μL) to a 5 mL Sephadex G25 gel filtration column with PBS as a mobile phase.

To functionalise the nanoparticles with FluPep ligand, this was incorporated into the initial ligand mix at the mole % indicated in the figure legends. However, the separation of free FluPep ligand (mwt 2967 Da) required, in addition to the final gel-filtration on Sephadex G25, six washes, each wash involving a 10-fold dilution of the nanoparticles on a 10 kDa cut off Nanosep filter.

The concentration of FluPep ligand in a nanoparticle preparation was estimated in two ways: for low mole % of FluPep ligand, where the majority of the nanoparticles are determined to be functionalised with a single FluPep ligand, the nanoparticle concentration was used as a proxy for FluPep ligand concentration, at higher mole % FluPep ligand, the number of peptidols incorporated into a 70:30 (mole/mole) mix matrix ligand shell of 8.8 nm gold nanoparticles (measured previously, 1200 peptidols, Duchense et al., 2008) was then multiplied by the mole % of FluPep ligand and by 1200.

#### Ion-exchange chromatography

Ion-exchange chromatography was performed on homemade mini columns of diethylaminoethyl (DEAE) and carboxymethyl (CM) Sepharose (GE Healthcare Bio-Sciences AB, Sweden). The chromatography gel slurry was packed into a white pipette tip (200 μL) using half the filter as a frit and equilibrated in PBS. Capped nanoparticles were concentrated and exchanged into the appropriate buffer using a a 10 kDa cut off Nanosep centrifugal filter. The nanoparticles were then applied to the column, the unbound fraction was recovered. Columns were washed with PBS and eluted with 1 M NaCl and then 2 M NaCl in 8 mM Na_2_HPO_4_, 15 mM KH_2_PO_4_, pH 7.4.

#### Ligand exchange assay

Purified ligand capped nanoparticles, 57 μL, 33 μL 10X PBS, and 10 μL DTT solutions at different concentrations (or milliQ water when the required concentration of DTT was 0) were added to a 384 well plate. A blank well, which only contained 100 μL of milliQ water, was used as reference. Spectra were acquired on duplicate wells at indicated times.

#### Calculation of the aggregation parameter (AP)

The surface plasmon absorption peak of 8.8 nm diameter gold nanoparticles is at 520 nm. When gold nanoparticles are aggregated, their surface plasmons couple causing a red shift in their plasmon absorbance to approximately 650 nm. The aggregation parameter (AP) was defined as (A _650 nm_-A_ref650nm_)/ (A_520nm_-A_ref520_), where A_650nm_ and A_520nm_ are the absorbance of gold nanoparticles at 650 nm and 520 nm, respectively, and A_ref650nm_ and A_ref520_ are the absorbance of water at 650 nm and 520 nm, respectively (Chen et al., 2012). For comparison of results, this primary stability parameter was normalised by dividing the AP value of control ligand capped gold nanoparticles measured in milli Q water where [DTT] = 0.

For silver nanoparticles diameter of approximately 10 nm, the surface plasmon absorption peak with a mix matrix ligand shell is at 410 nm. The AP for silver nanoparticle was defined as (A _600 nm_-A_ref600nm_)/ (A_410nm_-A_ref410_), where A_600nm_ and A_410nm_ are the absorbance of Ag nanoparticles at 600 nm and 410 nm, respectively and A_ref 600nm_ and A_ref 410_ are the absorbance of water at 600 nm and 410 nm, respectively.

#### Cell culture

Madin-Darby canine kidney epithelial cells [MDCK (ATCC CRL-2936)] were grown in Dulbecco’s modified Eagle’s medium (DMEM) supplemented with 5% (v/v) foetal calf serum (FCS) (Labtech International Ltd, East Sussex, UK), 1% (v/v) L-glutamine 200 mM, 1% (v/v) 100 U/mL penicillin and 1% (v/v) 100 μg/mL streptomycin and incubated in a humidified environment at 37°C under 5% (v/v) CO_2_ atmosphere. Cells were passaged with 0.05% (w/v) trypsin in the chelating agent, 1x Versene-EDTA (Gibco, Life Technologies, UK).

#### Preparation of influenza virus stock

MDCK cells were grown to 90% confluence in T25 tissue culture flasks, which corresponds to 7 × 10^6^ cells/flask. Then, the cell monolayer was washed with 2 × 5 mL PBS, and virus (A/WSN/33 H1N1 subtype) was added at a multiplicity of infection (MOI) of 0.001 in 2 mL DMEM made by diluting virus stock to give a 1000-fold final dilution. Cells were incubated with virus for 1 h at 37°C on a rocking platform. Virus containing medium was removed and the cell monolayer washed with 2 × 5 mL DMEM, then 5 mL N-acetyl trypsin (Sigma-Aldrich, Merck, UK), 2.5 μg/mL in DMEM, was added and incubated 24-48 hours at 37°C until significant cytopathic effect had developed to a point where the cells were lifting from the flask substrate. Medium was removed and centrifuged for 5 min at 2500 rpm to remove cell debris and the supernatant, which represented the viral stock, was stored at −80°C.

#### Virus plaque assay

MDCK cells were grown in 6-well plates (10^6^ cells/well) for two days. At confluence, monolayers of MDCK cells were then infected with a serial dilution of influenza virus inoculum (sufficient to obtain approximately 100 plaques per well) for 1 h at 37C° on a rocking platform. An agarose overlay was prepared by mixing equal volumes of 2 % (w/v) of pre-warmed (55°C) low melting agarose (Melford Laboratories Ltd, Blideston Road, Ipswich, UK) and the overlay solution [14 mL 10x MEM, 3.7 mL 7.5% (w/v) bovine serum albumin (fraction V, Sigma-Aldrich, UK), 1.4 mL L-glutamine, 2.6 mL 7.5 % (w/v) NaHCO_2_, 1.4 mL 1 M HEPES, 1.4 mL (1% (v/v) 100 U/mL penicillin and 1% (v/v) 100 μg/mL streptomycin), 44.8 mL H_2_O and 5μL N-acetyl Trypsin] to give a final 1 % (w/v) agarose mixture. After 1 h of incubation of the cells with the virus, the supernatant was removed from the plates and overlaid with 2-3 mL of the 1% (w/v) agarose overlay solution. The plates were left at room temperature for 15 minutes for the overlay to solidify and then inverted and placed in an incubator at 37 C°, 5% (v/v) CO_2_ for 3 days for plaques to develop. Cells were then fixed with 4 mL 10 % (v/v) neutral buffered formalin (Leica Biosystems Peterborough Ltd, Bretton Peterborough, Cambridgeshire) for 1 h, after which the formalin and overlay were removed and cells were stained with 0.1% (w/v in water) toluidine blue, rinsed in water, and left to dry before counting plaques.

## Results and Discussion

### Stability of FluPep functionalised gold nanoparticles

The mix matrix ligand shell of 70:30 mole/mole peptidol and alkanethiol ethylene glycol assembled on gold nanoparticles has well characterised stability with respect to ligand exchange and non-specific binding (Duchesne et al., 2008a, Chen et al., 2012), but the effect of incorporating the FluPep amino acid sequence at the C-terminus of the CVVVT matrix sequence is unknown. Since the molecular weight of FluPep ligand (2967 Da) is greater than that required for group separation on Sephadex G25 (1000 Da) additional purification by means of washes on a 10 kDa cut off Nanosep filter were included to ensure removal of any free FluPep ligand. When up to 5% (mole/mole) FluPep ligand was incorporated in the ligand matrix, the gold nanoparticles still eluted in the void volume of the Sephadex G25 column, so did not bind non-specifically to this chromatography matrix and their uv-vis spectrum in PBS was identical to that of control mix matrix gold nanoparticles (Fig. 1A). This indicates that the FluPep sequence did not reduce the stability of the gold nanoparticles under these standard conditions. A more stressful test is ligand exchange with small thiols (Chen et al., 2012, Duchesne et al., 2008a, Schulz et al., 2016, Schulz et al., 2013). Ligand-exchange results in a ligand shell that is unable to prevent electrolyte-induced aggregation of the nanoparticles, demonstrated by a decrease in the plasmon absorption at 520 nm.

**Fig. 1.**
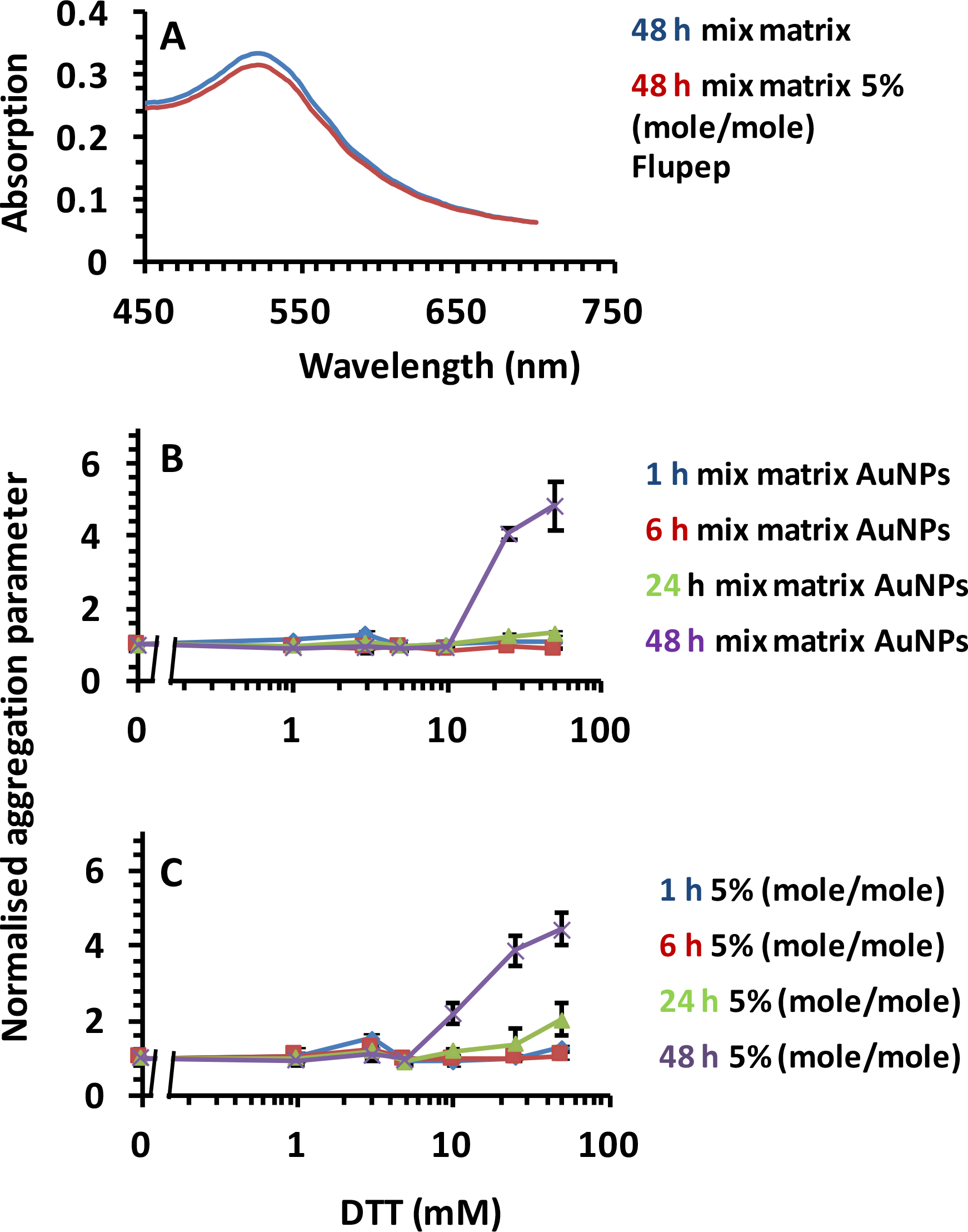
Stability of gold nanoparticles to DTT ligand exchange. (A) uv-vis spectra of mix matrix capped gold nanoparticles and mix matrix capped gold nanoparticles functionalised with 5% (mole/mole) Flu|Pep ligand in PBS. Time and dose-dependence of DTT ligand exchange for (B) mix matrix gold nanoparticles and for gold nanoparticles with a ligand shell incorporating 5% (mole/mole) FluPep ligand. Results in (b) and (C) are the mean ± SD of the calculated aggregation parameter (n=3).

Gold nanoparticles with a ligand shell incorporating 5% (mole/mole) FluPep ligand had a very similar resistance to ligand exchange with DTT as the control mix matrix protected gold nanoparticles. Thus, their aggregation parameter was unchanged up to 5 mM DTT, even after 48 h incubation (Figs 1 B, C). At 10 mM DTT after 48 h there was some evidence for ligand exchange, as the aggregation parameter was above 1.0 and at 25 mM DTT the ligand shell was clearly compromised. Nanoparticles incorporating lesser amounts of FluPep ligand (0.1% to 3 % mole/mole) were no less stable (Figs S1A-F). Consequently, the incorporation up to 5% (mole/mole) FluPep ligand in the ligand mixture did not reduce the stability of the gold nanoparticles with respect to ligand exchange and such nanoparticles could be used in cell culture medium.

### Purification of FluPep functionalised gold nanoparticles

To functionalise the nanoparticles, the peptide FluPep ligand was included in the ligand mix, its mole % in relation to the matrix ligand reflects its grafting density on the gold nanoparticles (Levy et al., 2006, Duchesne et al., 2008a, Duchesne et al., 2012, Free et al., 2012, Paramelle et al., 2015, Nieves et al., 2014). This can be determined by chromatography targeting specifically the grafted function, which also provides a means to purify the functionalised gold nanoparticles from those not functionalised, when the mole % of the functional ligand is low. Thus, when 10% of the functionalised gold nanoparticles bind to the chromatography column, most of these (95 %) will possess just one grafted functional ligand (Levy et al., 2006, Duchesne et al., 2008a). Since FluPep ligand, like FluPep, has a net charge at pH 7.4 of +6, cation-exchange chromatography was used to purify functionalised gold nanoparticles. Parallel chromatography was performed on the anion-exchanger DEAE-Sepharose to control for possible non-specific binding of FluPep ligand to Sepharose.

Mix matrix gold nanoparticles did not to CM Sepharose nor to DEAE Sepharose (Fig. S2), as described previously (Duchesne et al., 2008a). Similarly, when FluPep ligand was incorporated in the ligand shell there was no binding to DEAE Sepharose, indicating the absence of non-specific interactions with the chromatography resin (Fig. S2). In contrast, the FluPep functionalised gold nanoparticles bound to CM Sepharose and were eluted by increasing the electrolyte concentration of the eluent (Fig. 2). Thus, the FluPep functionalised gold nanoparticles ion-exchanged with a negatively charged resin, which is, therefore, suitable for their purification. Gold nanoparticles were synthesised with a range of mole % of FluPep ligand. After application of the gold nanoparticles to the column, the non-functionalised gold nanoparticles were collected in the flow through and the functionalised ones eluted. Quantification of the GNPs by UV-Vis spectrophotometry was used to estimate the relation of bound and unbound GNPs to the mole % of Flupep in the original ligand mixture. The data indicate that at 0.03 mole %, 10 % of the as-synthesised gold nanoparticles bound to the column and thus statistically most (~95 %) of these gold nanoparticles will possess just a single FluPep ligand on their surface (Levy et al., 2006). At higher mole % the number of FluPep ligands per nanoparticle will increase. It is interesting to note that not all gold nanoparticles were observed to bind to the CM Sepharose resin at higher mole % of FluPep ligand, something that has been observed previously with other functional peptides (Nieves et al., 2014, Paramelle et al., 2015). Nonetheless, this straightforward purification of the functionalised gold nanoparticles means that the effects on influenza virus infectivity of mono-*versus* plurifunctionalisation can be determined. Moreover, since it is known that gold nanoparticles passivated with the mix matrix (70:30 molar ratio of SH-(CH_2_)_11_EG_4_ and CVVVT-ol) have an average of 1200 peptidols per nanoparticle (Duchesne et al., 2008a), it is possible to estimate the average number of FluPep ligands grafted on the gold nanoparticles by simply using their mole % in the ligand mixture.

**Fig. 2.**
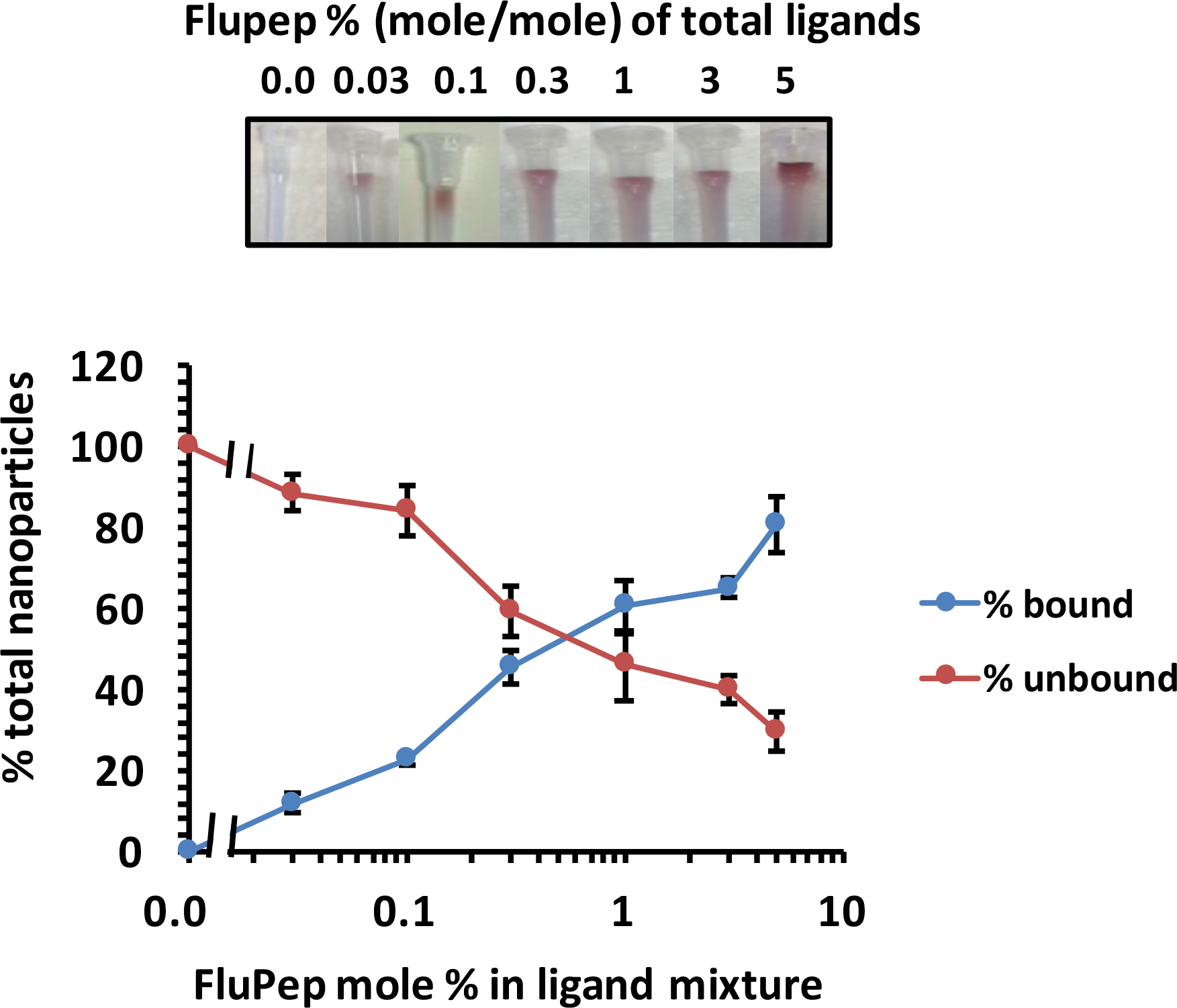
Purification of FluPep ligand functionalised gold nanoparticles by CM-Sepharose cation-exchange chromatography. Gold nanoparticles functionalised with different mole % FluPep ligand were subjected to chromatography on CM-Sepharose. Top, images of columns after loading and washing with PBS. Bottom, quantification of unbound (through and PBS wash fractions) and bound (eluted with 2 M NaCl) fractions. Results are the mean ± SD (n=3).

### Anti-flu activity of FluPep and FluPep ligand

The MDCK are susceptible to both influenza type A and B viruses and are routinely used to measure ‘flu virus infectivity. The principle is that following virus binding, virus will infect a cell, reproduce and lyse that cell; released virus then infects neighbouring cells, the overlay preventing long-range diffusion of virus. After three days, the area of lysed cells is visualised as a plaque that does not stain with toluidine blue. In control (no virus) and vehicle (DMSO) treated cells, the cell monolayer in the well was evenly stained (Fig. S3). In the presence of virus, there were substantial areas where cells had lysed and there was no staining. Virus titre was adjusted so that the cleared lysed cell areas corresponded to individual plaques, i.e., clear circles that could be distinguished from one another and so counted (corresponding to ~100 plaques/well). When FluPep was added with virus there was a reduction in the number of plaques and this was concentration-dependent (Fig 4, A). Counting plaques in multiple experiments allowed the determination of the dose-response and the IC_50_. In these experiments the IC_50_ of the original FluPep sequence was found to be of the order of 140 pM (Fig. 3). This is less potent than described in the original publication, where an IC_50_ of 14 pM was measured. The source of this discrepancy is unknown, but may relate to differences in cells such as passage number, and/or virus preparations. The FluPep ligand used to functionalise gold nanoparticles was slightly less potent, with an IC_50_ of around 210 pM, suggesting that the N-terminal extension CVVVTAA reduced the antiviral activity (Fig 3 and Table 1).

**Fig. 3.**
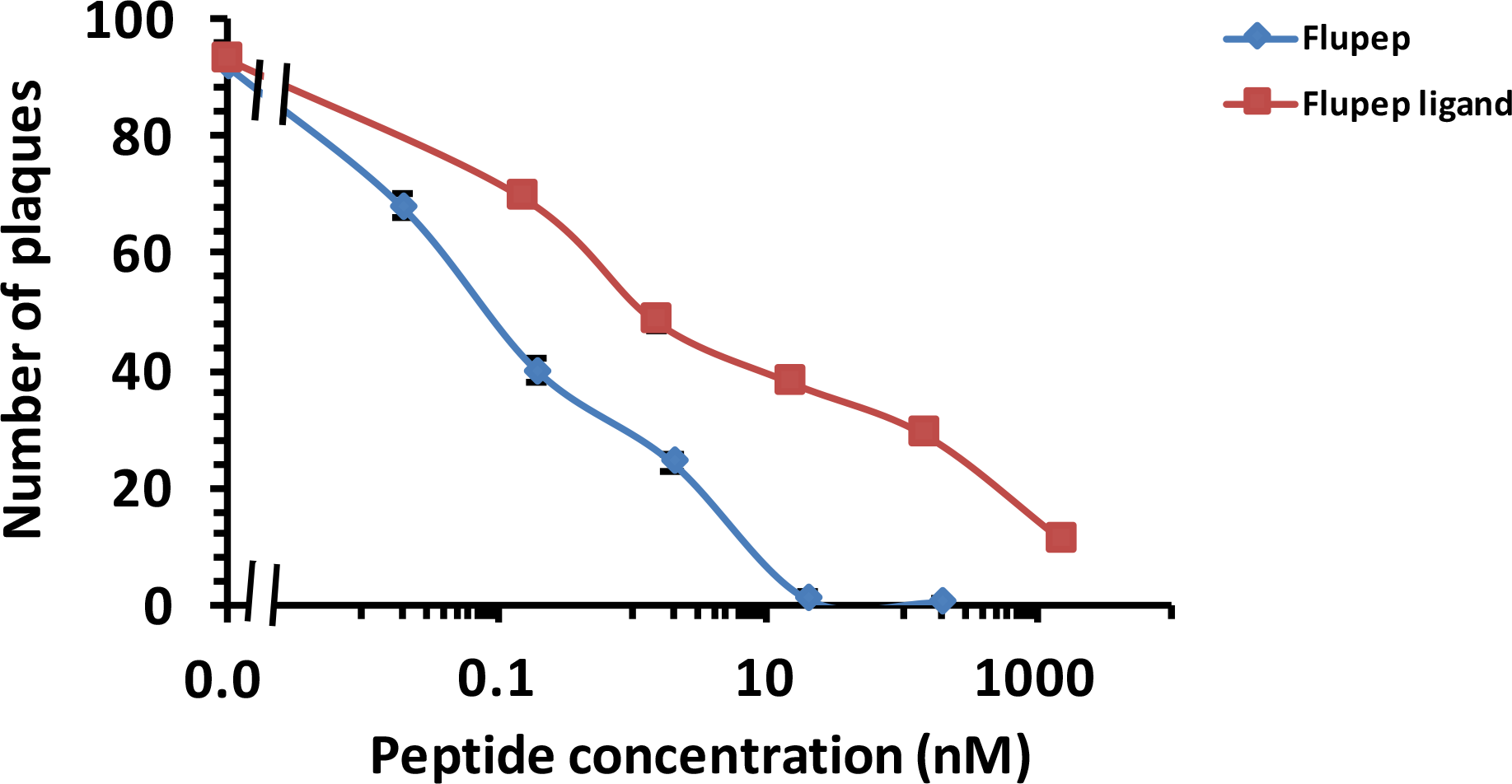
Determination of the half maximal inhibitory concentration (IC_50_) of FluPep and and FluPep ligand in a plaque assay. Virus and peptides were added to MDCK cell monolayers, and a plaque assay performed, as described in Materials and Methods. Results are the mean ±SD (n=3).

**Table 1:**
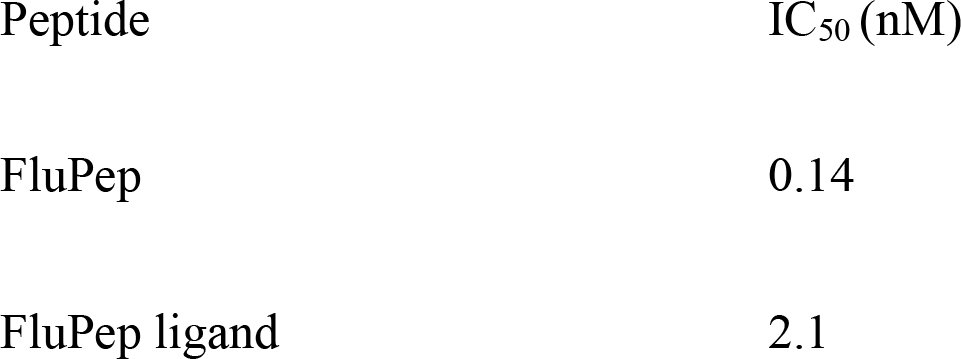
IC_50_ of viral plaque formation by FluPep and FluPep ligand.

### Anti-flu activity of FluPep functionalised gold nanoparticles

The addition of control mix matrix passivated nanoparticles along with the influenza virus did not change the number of plaques (Fig. 4A). Thus, these nanoparticles had no effect on viral infectivity. However, when purified mix matrix capped gold nanoparticles functionalised with 5 % (mole/mole) FluPep ligand were added with virus, there was a marked decrease in the number of plaques, i.e. reduced virus infectivity (Fig. 4A). Thus, FluPep ligand’s antiviral activity was maintained when it was conjugated to gold nanoparticles.

**Fig. 4.**
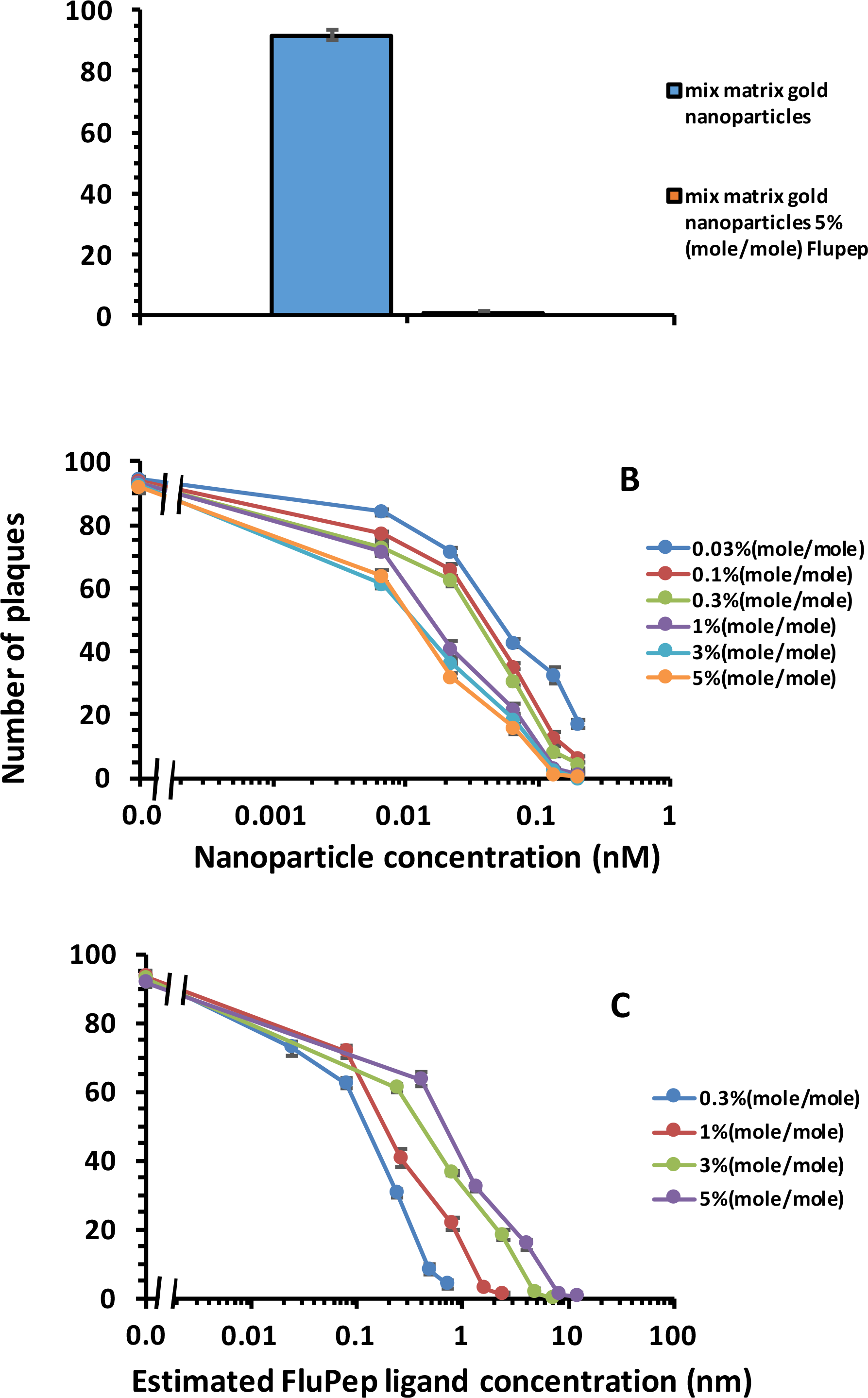
Effect of gold nanoparticles functionalised with FluPep ligand on influenza virus plaque formation. Virus and gold nanoparticles functionalised with FluPep ligand at the indicated mole % were added to MDCK cell monolayers and a plaque assay performed. (A) Number of plaques observed in the presence of mix matrix control gold nanoparticles and gold nanoparticles functionalised with 5 % (mole/mole) FluPep ligand. (B) Dependence of inhibition of plaque formation on the concentration of gold nanoparticles and the % (mole/mole) of FluPep ligand. (C) Same data as in (B), but expressed in terms of the estimated FluPep ligand concentration. Results and the mean ±SD (n=3).

For gold nanoparticles functionalised with 0.03% (mole/mole) FluPep ligand, the number of plaques started to decrease at 20 pM gold nanoparticles and reached a minimum of approximately two to eight plaques at 200 pM (Fig. 4B). As the grafting density of FluPep ligand was increased, so did the antiviral activity of the FluPep functionalised gold nanoparticles, to reach a maximum at 5 % (mole/mole) FluPep ligand (Fig. 4B). This is reflected by the decreased IC_50_, which shows a 4.5-fold greater potency of nanoparticles functionalised with 5 % (mole/mole) FluPep ligand, when compared to nanoparticles functionalised with just 0.03 % (mole/mol) FluPep ligand (Table 2). The nanoparticles were always purified by cation-exchange chromatography (Fig. 2), so in all cases nanoparticles had at least one FluPep ligand. When most nanoparticles have just a single FluPep ligand (0.03 % and 0.1 % mole/mole, Fig. 2), the nanoparticle concentration is a reasonable proxy for FluPep ligand concentration. Taken together, these data indicate that the potency of FluPep ligand on the nanoparticles is greater than that of free FluPep and of the native FluPep peptide (Tables 1 and 2).

An interesting question is whether the potency of the FluPep functionalised nanoparticles will increase as the FluPep grafting density increases, or whether there would be an optimal FluPep ligand density on the nanoparticle’s surface. At higher mole % FluPep ligand (0.3 % mole/mole and above) many, if not all nanoparticles, will be functionalised with two or more FluPep ligands. In these cases, the nanoparticle concentration is no longer a proxy for the FluPep ligand concentration. Therefore, to estimate the concentration of FluPep ligand in these nanoparticles, we used the measured number of peptidols in the mix matrix ligand shell (1200, Duchesne et al., 2008a) and the mole % of FluPep ligand. Using these estimates, it was apparent that the potency of the FluPep-nanoparticle conjugates declined as the number of FluPep ligands per nanoparticle increased (Fig 4 C). This is reflected in the IC_50_ of the nanoparticles, calculated on the basis of FluPep ligand concentration (Table 2 for 0.03 % and 0.1 % mole/mole FluPep ligand and Table 3 for ≥0.3 % mole/mole FluPep ligand). It would appear then that the FluPep ligand nanoparticle conjugate is no more potent than the free peptides from between 0.3% and 1% (mole/mole) FluPep ligand (Tables 1-3). The difference in IC_50_ between nanoparticles with a single FluPep ligand (0.03 % and 0.1 % mole/mole FluPep ligand) and nanoparticles with the highest grafting density (5 % mole/mole FluPep ligand) is ~10-fold (Tables 2 and 3). This is likely greater than any uncertainty in the estimation of concentration of FluPep ligand, which can be gauged from the departure of the data in (Fig. 2) from the ideal, which was described previously (Levy et al., 2006). Therefore, these data suggest that as the number of FluPep ligands on the nanoparticle surface increases to two or more, they are less active. However, the data also demonstrate that free FluPep and FluPep ligand are less active than nanoparticles grafted with a single FluPep ligand. One explanation for this is that monomeric FluPep is the active species, but the peptide may self-associate to produce dimers/oligomers with lower activity with respect to inhibiting influenza virus infectivity. The hydrophobicity of the FluPep sequence could be an important factor in such self-association. Moreover, self-association of the FluPep sequence might be greater for FluPep on a nanoparticle surface. For example, the association of functional peptides grafted into a peptide ligand shell on nanoparticles with themselves has been observed directly previously (Duchesne et al., 2008b). Alternatively, the targets for FluPep may be separated by a distance similar or greater than their average physical separation on the nanoparticle surface, resulting in only some of the grafted FluPep ligands on the nanoparticle surface being able to exert an antiviral activity.

**Table 2.:**
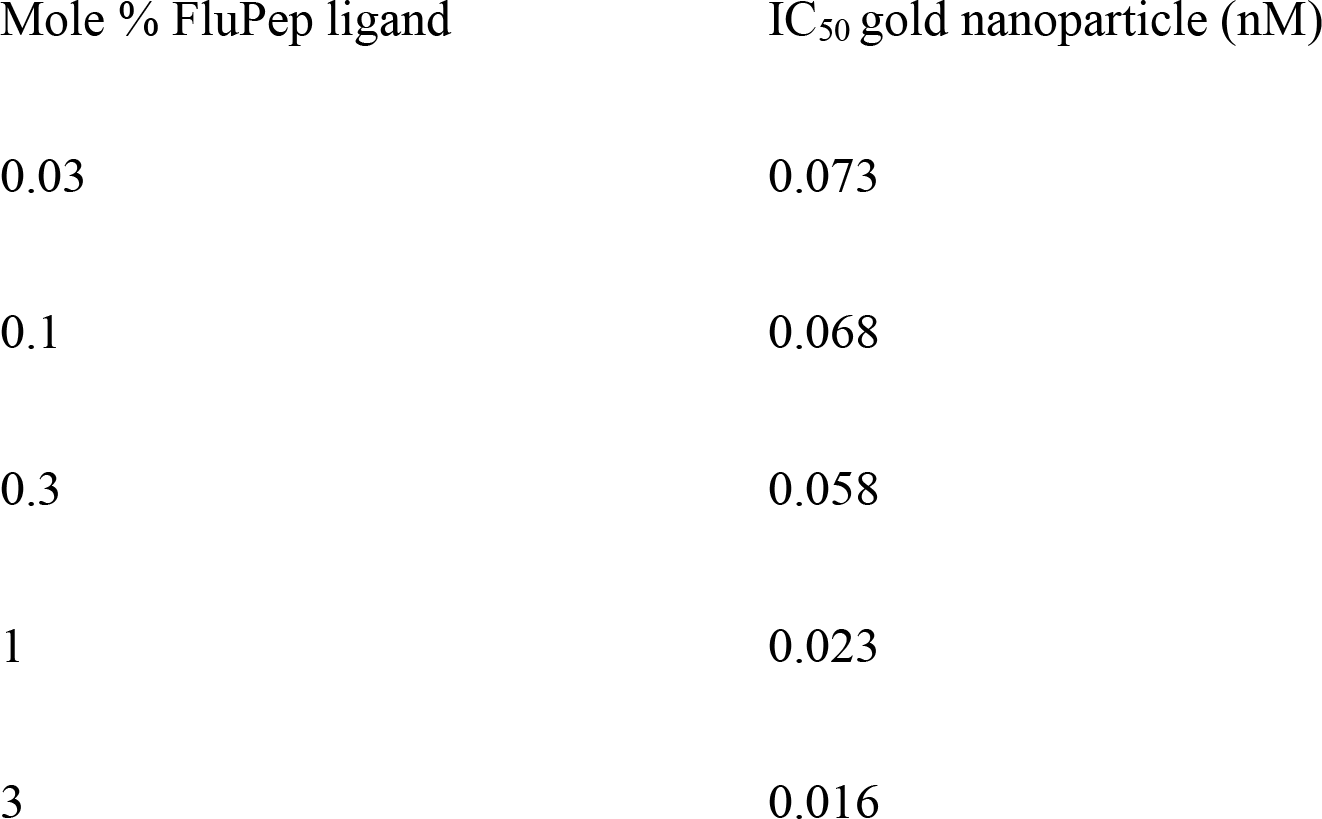
IC_50_ of viral plaque formation by golf nanoparticles functionalised with different mole % of FluPep ligand, calculated with respect to gold nanoparticle concentration.

**Table 3:**
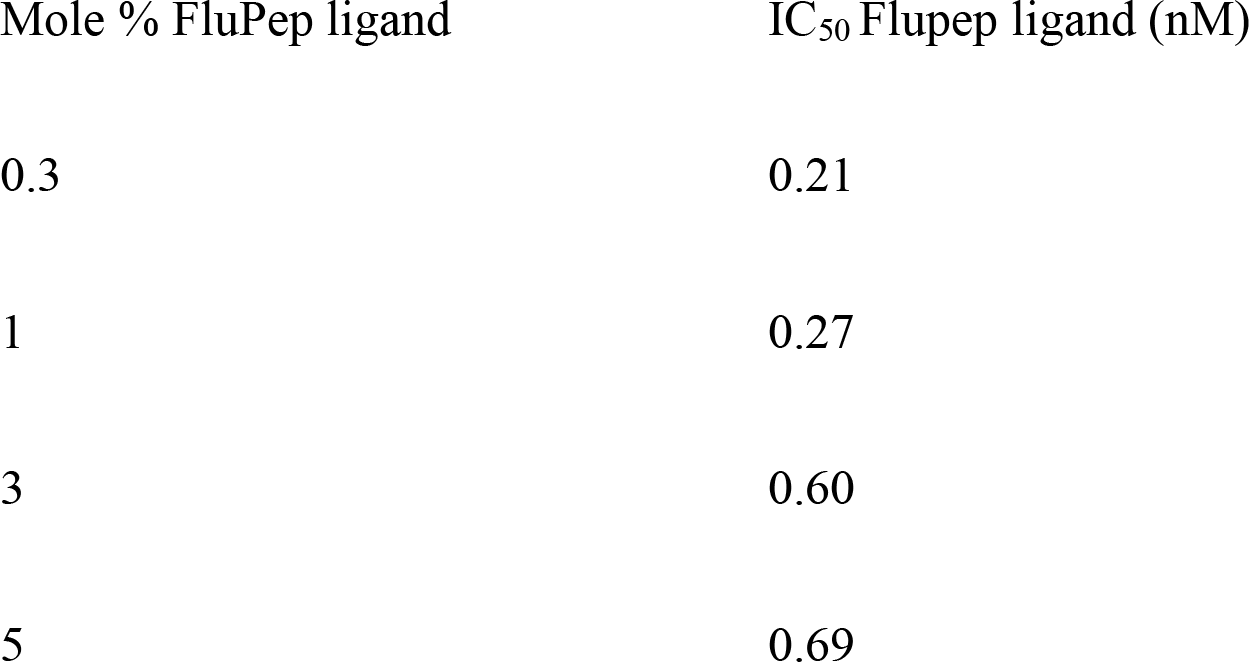
IC_50_ of viral plaque formation by gold nanoparticles functionalised with different mole % FluPep ligand calculated with respect to the estimated concentration of FluPep ligand.

| Mole % FluPep ligand | IC<sub>50</sub> Flupep ligand (nM) |
| --- | --- |
| 0.3 | 0.21 |
| 1 | 0.27 |
| 3 | 0.60 |
| 5 | 0.69 |

### Stability of FluPep functionalised silver nanoparticles

Silver has well-established anti-microbial properties. It was of interest to determine whether FluPep functionalised silver nanoparticles possessed a similar anti-flu activity to their gold counterparts. The mix matrix ligand shell has been shown to impart good stability to silver nanoparticles (Free et al., 2012). First, the effect of incorporating FluPep into the ligand shell on nanoparticle stability was measured. Similarly to gold nanoparticles, up to 5% (mole/mole) FluPep ligand incorporated into the ligand matrix had no discernible effect on the handling and purification of the silver nanoparticles: the silver nanoparticles did not bind non-specifically to Sephadex G25, as they eluted in the void volume and their uv-vis spectrum in PBS was indistinguishable from that of control mix matrix silver nanoparticles (Fig. 5A). When challenged by a small thiol, DTT, the silver nanoparticles passivated by the mix matrix ligand shell were somewhat more prone to ligand exchange than their gold counterparts. Thus, after 3 h and 6 h in 50 mM DTT a small increase in aggregation parameter was apparent (Fig. 5B). After 24 h and 48 h, the aggregation parameter started to rise at 3 mM DTT, which was most pronounced at 48 h. Importantly, the inclusion of 5% (mole/mole) FluPep ligand in their ligand shell did not change the stability of the silver nanoparticles with respect to DTT-mediated ligand exchange (Fig. 5C).

**Fig. 5.**
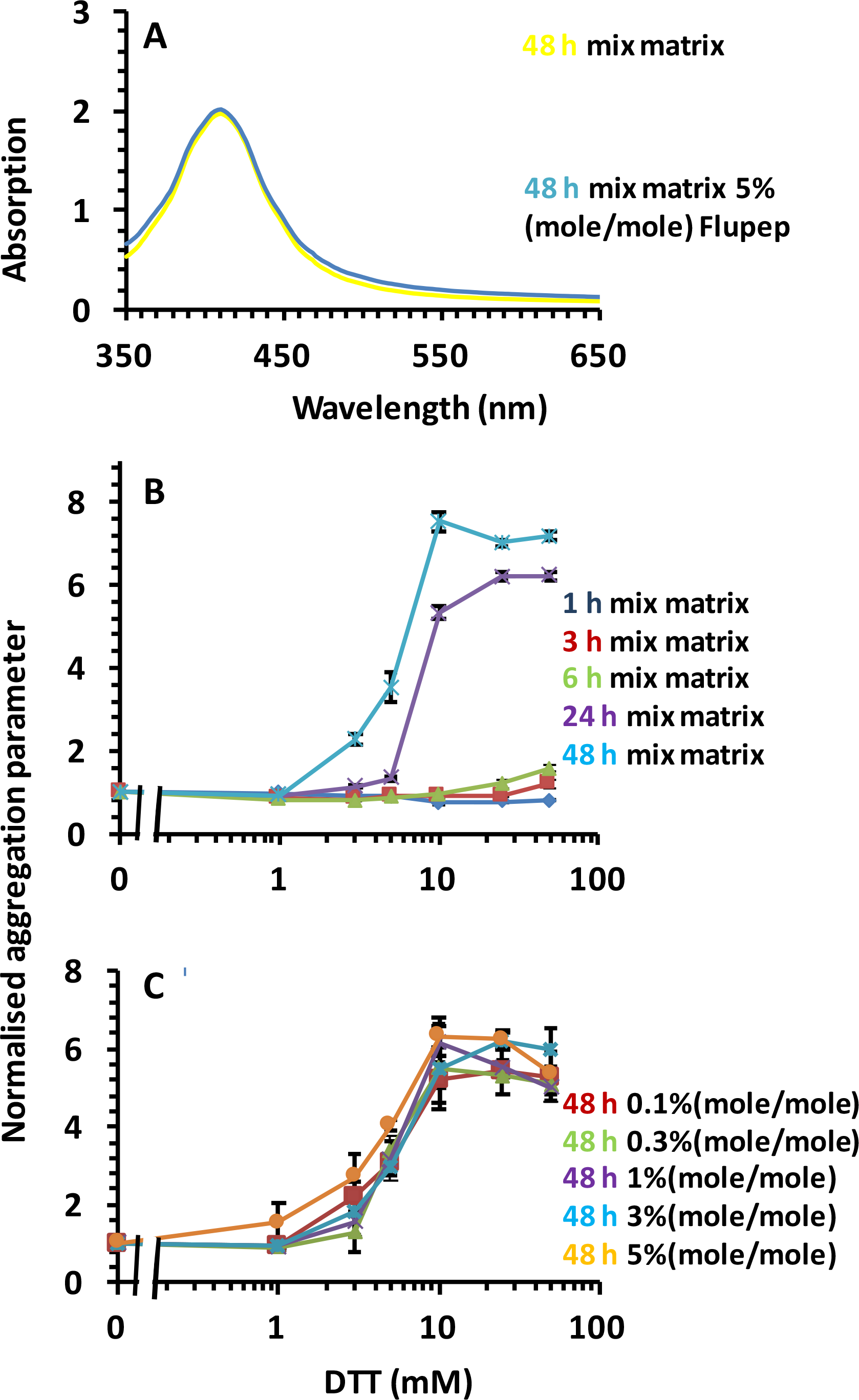
Stability of silver nanoparticles to DTT ligand exchange. (A) uv-vis spectra of mix matrix capped silver nanoparticles and mix matrix capped silver nanoparticles incorporating 5% (mole/mole) Flu|Pep ligand in PBS. Time and dose-dependence of DTT ligand exchange for (B) mix matrix silver nanoparticles and (C) silver nanoparticles incorporating different mole % FluPep ligand. Results are the mean calculated aggregation parameter ± SD (n=3).

Purification of the FluPep functionalised silver nanoparticles was achieved by cation-exchange chromatography on CM-Sepharose. As the percentage of FluPep incorporated into the ligand shell increased, so did the percentage of nanoparticles bound to CM-Sepharose (Fig. 6). This suggests that the mole % FluPep ligand in the initial mixture of ligands added to the nanoparticles reflects its incorporation into the ligand shell (Levy et al., 2006, Duchesne et al., 2008a). Thus, similarly to gold nanoparticles, when 10 % of the total nanoparticle preparation bound to the CM-Sepharose column, ~95% of the bound nanoparticles will possess a single FluPep ligand (Levy et al., 2006); at higher mole % of FluPep ligand and increasing proportion of the silver nanoparticles will incorporate more than one FluPep ligand.

**Fig. 6.**
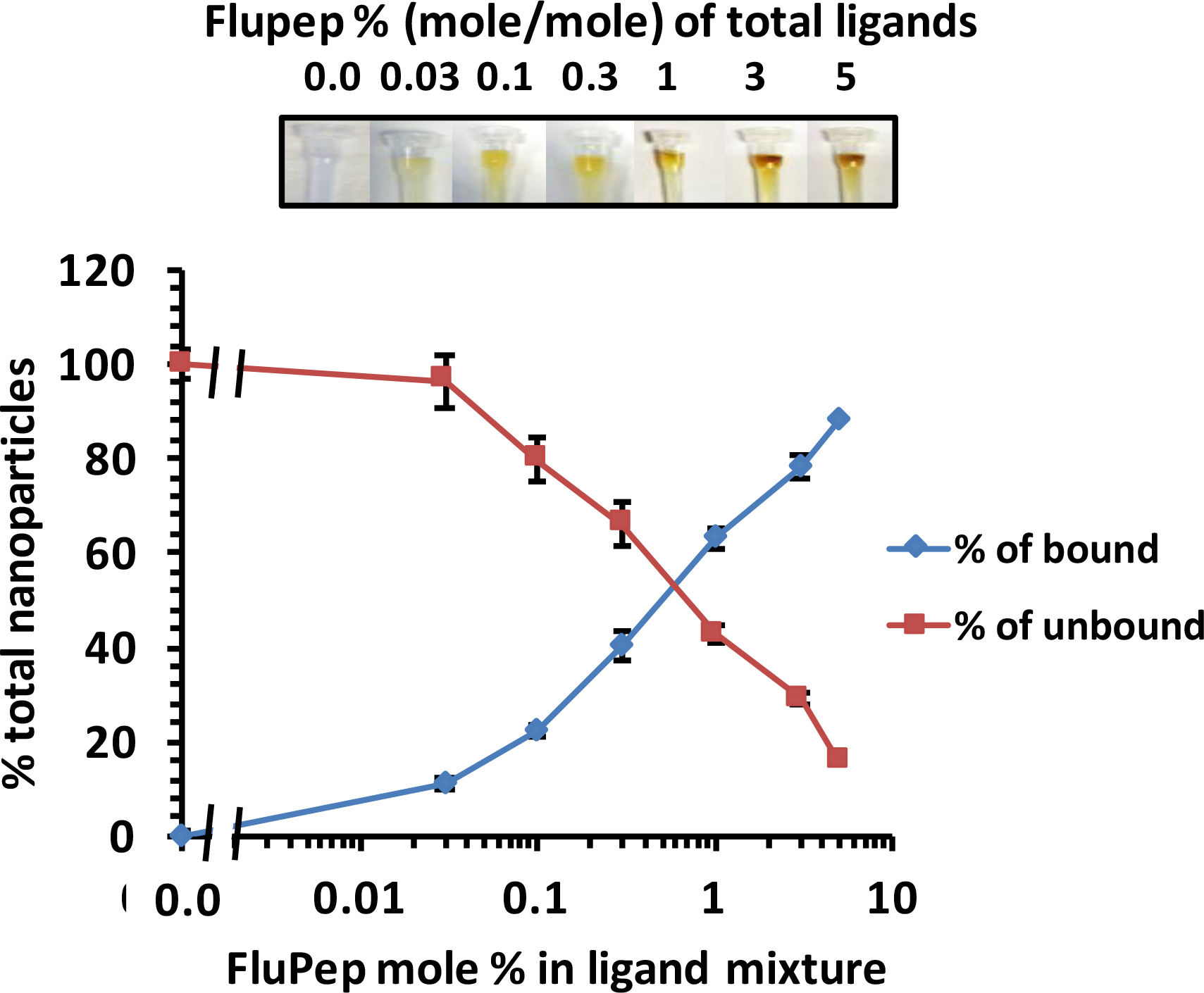
Purification of FluPep ligand functionalised silver nanoparticles by CM-Sepharose cation-exchange chromatography. Silver nanoparticles functionalised with different mole % FluPep ligand were subjected to chromatography on CM-Sepharose. Top, images of columns after loading and washing with PBS. Bottom, quantification of unbound (through and PBS wash fractions) and bound (eluted with 2 M NaCl) fractions. Results are the mean ± SD (n=3).

### Anti-flu activity of FluPep ligand incorporated to silver nanoparticles

Control mix matrix passivated silver nanoparticles had no effect on viral infectivity (Fig. 7). It is the silver (Ag^+^) ions that exert anti-microbial activity (Hsueh et al., 2015). Thus, this result indicates that there is not a substantial release of Ag^+^ ions from the silver nanoparticles during the experiment. This concurs well with our data (Fig. 5) and previously reported observations (Free et al., 2012) demonstrating that the mix matrix ligand shell imparts good stability to the silver nanoparticles. Incorporation of FluPep ligand into the mix matrix ligand shell caused a marked reduction in number of plaques (Fig. 7A). As the mole % of FluPep ligand in the ligand shell increased, so did the antiviral activity of these particles (Fig. 7A and Table 4). Around 10% of silver nanoparticles functionalised with 0.03 % (mole/mole) FluPep ligand bind to the CM-Sepharose column, indicating that the majority of these will carry just a single FluPep ligand (Fig. 6), so the nanoparticle concentration will equate to the FluPep ligand concentration. These nanoparticles are as potent as free FluPep and more potent than free FluPep ligand (Tables 1 and 4). However, FluPep ligand functionalised silver nanoparticles were less effective at inhibiting influenza virus infectivity than the corresponding gold nanoparticles (Tables 2 and 4). At higher mole % FluPep ligand (≥0.1 % mole/mole) many, if not all nanoparticles, will be functionalised with two or more FluPep ligands. As for gold nanoparticles, the nanoparticle concentration will no longer equate to the FluPep ligand concentration. To estimate the concentration of FluPep ligand in these nanoparticles, we assumed the same grafting density of peptidols on silver nanoparticles as for gold nanoparticles (1200 peptidols/nanoparticle, Duchesne et al., 2008a) to calculate an estimated FluPep ligand concentration (Figs 7B and Table 5). Using this estimate of FluPep ligand concentration, similar to gold nanoparticles, the data indicate that the relative potency of FluPep ligand decreases as its grafting density on the nanoparticles increases.

**Fig. 7:**
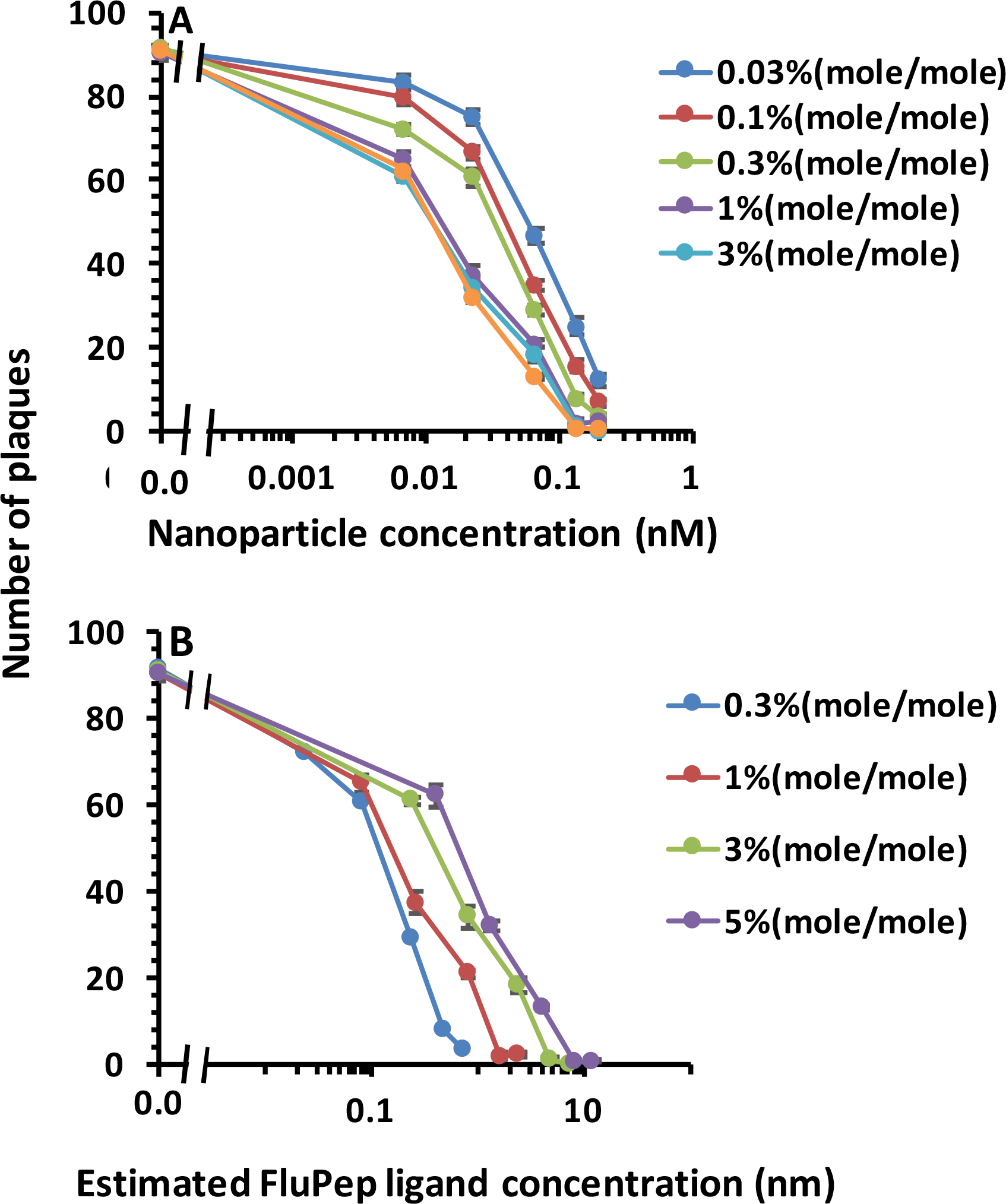
Determination of the half maximal inhibitory concentration (IC_50_) of silver nanoparticles functionalised with FluPep ligand. Virus and silver nanoparticles functionalised with FluPep ligand at the indicated mole % were added to MDCK cell monolayers and a plaque assay performed. (A) Dependence of inhibition of plaque formation on the concentration of silver nanoparticles and the % (mole/mole) of FluPep ligand. (C) Same data as in (B), but expressed in terms of estimated FluPep ligand concentration. Results and the mean ±SD (n=3).

**Table 4:** IC_50_ of viral plaque formation by silver nanoparticles functionalised with different mole % FluPep ligand.

| Mole % FluPep ligand | IC<sub>50</sub> silver nanoparticles (nM) |
| --- | --- |
| 0.03 | 0.14 |
| 0.1 | 0.077 |
| 0.3 | 0.056 |
| 1 | 0.019 |
| 3 | 0.016 |
| 5 | 0.015 |

**Table 5:** IC_50_ of viral plaque formation by silver nanoparticles functionalised with different mole % FluPep ligand, calculated with respect to the estimated concentration of FluPep ligand.

| Mole % FluPep ligand | IC<sub>50</sub> FluPep ligand (nM) |
| --- | --- |
| 0.3 | 0.20 |
| 1 | 0.24 |
| 3 | 0.60 |
| 5 | 0.68 |

## Conclusion

Influenza viruses pose a substantial danger to global health, because these viruses undergo rapid mutations in their genomes, they transmit with ease *via* the respiratory route, and there are substantial animal reservoirs. Currently, the preventative and remedial therapies against these viruses are inadequate. In the current study, we have functionalised nanoparticles with FluPep, a peptide demonstrated to inhibit influenza A virus subtypes, including H1N1, H3N2 and H5N1(Nicol et al., 2012). Monofunctionalised nanoparticles were more active from the perspective of FluPep ligand concentration than plurifunctionalised nanoparticles or free peptide. These data highlight that a nanoparticle formulation of such peptide inhibitors of ‘flu virus may be particularly effective. Purification of the functionalised nanoparticles by cation-exchange chromatography indicated that the FluPep ligand sequence was exposed to solvent, rather than associated with the mix matrix ligand shell, a conclusion supported by the observation that the gold and silver nanoparticle FluPep ligand conjugates possessed antiviral activity. At low stoichiometry of FluPep ligand: nanoparticle conjugation, the potency of the FluPep ligand is enhanced compared to that of the free peptide. The FluPep amino acid sequence is hydrophobic and its solubilisation requires dimethyl sulfoxide (DMSO) to be solubilised. Although a useful solvent, its use in therapies is problematic due to DMSO’s potential adverse reactions in some individuals such as a sensation of burning, vesiculation, dryness of skin and local allergic reactions (Cavaletti et al., 2000, Delatorre et al., 1981, Worthley and Schott, 1969). Thus, conjugation to nanoparticles may provide a means to deliver effectively and safely FluPep ligand with enhanced activity in a solvent-free formulation. Though the mix matrix ligand shell prevents the dissolution of the silver, its modification will reduce its effectiveness (Free et al., 2012). Thus, it will be possible to design silver nanoparticles that act through FluPep ligand and released Ag^+^ ions. The pulmonary route to deliver drugs against respiratory infections is well established, therefore, delivery of functionalised nanoparticles against ‘flu viruses is highly feasible. In conclusion, nanoparticle formation of FluPep ligand or analogous peptides offers a route to new treatments for ‘flu and other respiratory pathogens.

## Acknowledgements

ZA was supported by a PhD studentship award from the Iraqi Ministry of Higher Education. DGF acknowledges the support of North West Cancer Research. The authors thank Dr David Paramelle, Institute of Materials Research and Engineering, Singapore, for critical reading of the manuscript.

